# Fibronectin Tethering as a Shear-thinning Bio-Annealing Strategy for Granular Hydrogels

**DOI:** 10.64898/2026.10.20.760419

**Authors:** Ethan J. Nicklow, Olivia D. Ernst, Chris Highley, Donald R. Griffin

## Abstract

Granular hydrogel scaffolds (GHS) are injectable, integrative biomaterials built from hydrogel microparticles that permit rapid cell infiltration independent of scaffold degradation. Microporous annealed particle (MAP) scaffolds, a sub-class of GHS, use inter-microparticle crosslinks to anneal adjacent particles, providing greater mechanical stability than purely jammed assemblies. A growing range of inter-particle crosslinking strategies—including covalent photo-initiated radical polymerization and reversible guest-host chemistries, some intended to mimic extracellular matrix (ECM) viscoelasticity—focus on mechanical stabilization, while surface bioactivity is typically added separately via orthogonally tethered cell-adhesion ligands such as RGD, a peptide derived from fibronectin (FN), a key mediator of cell-ECM and ECM-ECM interactions. Here, we present an alternative that unifies these functions: coating MAP microparticle building blocks with covalently bound fibronectin to create fibronectin-modified MAP (FN-MAP). FN-MAP exhibits shear-thinning annealing with complete recovery of scaffold mechanics after physical mixing. Human lung fibroblasts respond favorably, adopting spread morphologies with high viability. Translationally, FN-MAP shows proof-of-principle promise for 3D bioprinting (greater stability and higher print resolution) and wound healing (increased epidermal closure in a diabetic mouse model). FN-MAP thus delivers both increased scaffold strength and bioactivity without exogenous soluble linkers or mixed particle populations.

**Table of Contents:** Porous scaffolds encourage tissue regeneration but typically need separate components to be both strong and bioactive. By coating granular hydrogel scaffolds with fibronectin, a natural protein central to wound repair, we simultaneously boost material strength and cell-instructive signaling. This single-protein strategy produces stronger, more printable scaffolds, promotes healthy cell activity, and enhances wound closure in a challenging diabetic mouse model.

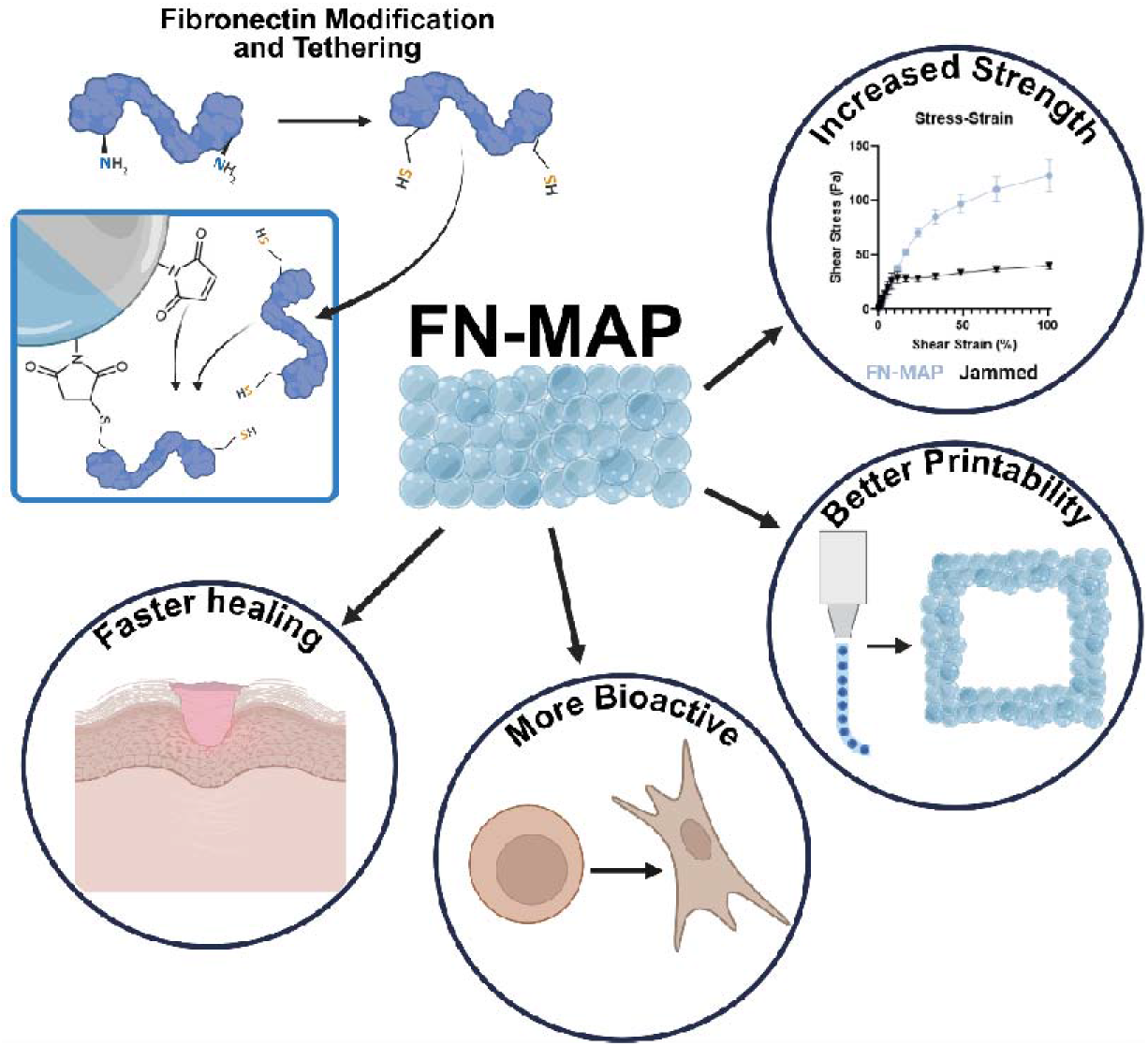

## 1. Introduction

Granular hydrogels are a biomaterial composed of packed hydrogel microparticles that form a porous, interconnected interstitial space between individual microparticles.^[1,2]^ This open architecture facilitates rapid molecular diffusion, accelerated cell infiltration and growth, and even the formation of complex tissues (e.g., vasculature, muscle fibers).^[3–5]^ The packed microparticles exhibit shear-thinning behavior under applied stress, enabling minimally invasive delivery via injection without disrupting the internal scaffold architecture^[6]^. Like bulk hydrogels, granular hydrogels can be fabricated from a variety of polymers and exhibit tunable bioactive and mechanical properties. Further, by incorporating a secondary mechanism to crosslink adjacent microparticles, referred to as annealing, the granular assembly is stabilized into a cohesive microporous annealed particle (MAP) scaffold that provides enhanced mechanical integrity. MAP scaffolds have shown promise as both a long-term 3D cell culture platform^[7]^ and as a clinically applicable scaffold for wound healing, stroke, cartilage repair, and tissue augmentation.^[4,8–11]^.

One key factor in these granular systems is the nature of particle-particle interactions; purely jammed systems rely primarily on friction for stability,^[6]^ while others, like MAP, rely on designed chemical crosslinks between particles.^[4,12]^ As the viscoelastic nature of endogenous extracellular matrix (ECM) environment has become better understood, MAP systems have been developed to leverage non-covalent crosslinking that better mimics the mechanical properties of native tissue to improve cell-material interactions.^[13,14]^ Stabilizing these scaffolds is particularly relevant as they must withstand ranges of physiological stress and strain when implanted in tissue or applied to wounds; bulk hydrogels, while lacking the ability for rapid cell infiltration, can offer stability at the cost of fibrotic responses *in vivo*.^[15,16]^ Recent work has focused on the strengthening of MAP scaffolds through cellular integration and matrix deposition, but synthetic scaffolds still struggle to achieve integrity that natural ECM embodies within strong, but dynamic materials.^[7]^ While the first MAP scaffold annealing chemistry relied on harnessing endogenous Factor XIIIa, a cell-compatible transglutaminase, to anneal adjacent particles via custom oligopeptide moieties, this approach was irreversible and slow. Subsequent iterations of MAP scaffolds have incorporated light-mediated covalent crosslinking that, while still irreversible, provided vastly increased temporal and spatial control.^[4,15,17]^ More recently, shear-thinning strategies to strengthen scaffolds via reversible bonds between particles have been reported (i.e., guest-host chemistries ^[18–21]^). These published systems harness heterogeneous particle populations^[18,22]^ or soluble linkers introduced via the open pores.^[23]^ Both of these approaches potentially limit ease-of-use and predictability in clinical settings, as incomplete mixing of particle populations or insufficient diffusion of soluble linkers may compromise scaffold stability, respectively. This manuscript explores the use of a homogeneous system with reversible behavior (i.e., shear thinning) by drawing inspiration from physiological protein interactions to simplify the design for better ease-of-use and predictability.

We were struck by previous results demonstrating an ordered ECM deposition in the interstitial space of synthetic MAP scaffolds, with fibronectin (FN) assembling at the material surface and seemingly serving as an intermediary to a stable collagenous ECM^[24]^. FN serves a particularly important role in physiological ECM assembly, organizing and maintaining connections through the ECM and playing a crucial role in multiple stages of tissue repair and regeneration while helping to orchestrate the formation of new tissue.^[25–28]^ It is an abundant glycoprotein found in plasma and throughout the ECM, acts as a potent mediator of cell-matrix interactions and forms fibrils that connect and support the network of ECM structural proteins.^[29]^ Cell-secreted fibronectin (FN) assembles into fibrils via cryptic FN-FN binding sites that change conformation with mechanical forces to allow for variable protein and cell binding. Plasma FN is secreted by hepatocytes and exists as a soluble dimer until it is transported to tissues or interacts with platelets and fibrin in a clot.^[29,30]^ It has a variety of well-characterized domains for interacting with other ECM proteins (i.e., collagen), cell surface integrins, and other ECM components. One significant cell-binding domain contains the RGD (Arg-Gly-Asp) sequence, a widely used cell-adhesion ligand attached to synthetic hydrogels to increase their bioactivity.^[31]^ Although other ligands have been explored (i.e., laminin-derived IKVAV and elastin-derived VPVAPG), RGD remains the most widely used.^[31,32]^ Beyond attaching RGD, groups have explored the incorporation of full-length FN via PEGylation and saw benefits in a bulk hydrogel model.^[33]^ The spatial segregation of FN in ECM developed within MAP suggests that FN can extend its structural role in ECM assembly into nascent ECM within synthetic scaffolds^[7,24]^. Based on this observation, we hypothesized that, by intentionally coating microparticles with FN, we could harness its ability to stabilize assembling ECM. Here, we investigate attachment of full-length FN to hydrogel microparticles to form bioactive MAP scaffolds stabilized via fibronectin binding interactions. We hypothesized fibronectin attachment would confer increased stability through ECM-derived annealing interactions that are reversible and self-healing while retaining enhanced scaffold bioactivity.

To introduce ECM-annealed MAP, we attached fibronectin to hydrogel microparticles after synthesis to allow for more natural presentation of FN than incorporation into the polymer network itself, preserving more bioactivity and avoiding complications with gelation. We subsequently demonstrated that scaffolds composed of fibronectin-coated particles (FN-MAP) do have increased strength and yield strain compared to purely packed systems, and that this increased stability is recoverable after repeated high strain. With these beneficial mechanical properties, we probed the bioactive effects of FN-MAP, observing increased fibroblast spreading and morphology while ensuring high viability is maintained. We also investigated the translational use of FN-MAP as an *in vitro* bioink and *in vivo* for wound healing. In these functional outcomes, we found that FN-MAP facilitates high print resolution and faster epidermal closure in a murine diabetic wound model, indicating significant promise even in complex wound environments. Altogether, we present evidence that FN-MAP offers predictable bioactivity and self-healing stability in a homogenous, injectable scaffold with applications in tissue formation and regeneration.

## 2. Results and Discussion

### 2.1 Characterization of fibronectin modification and attachment to MAP scaffolds

Although fibronectin has been incorporated into or coated on to bulk hydrogels,^[33]^ it has not been attached to granular hydrogels. As a platform for fibronectin presentation, fully PEG backbone-based particles (i.e., without peptide linkers) with excess maleimide groups for downstream attachment can be consistently synthesized with replicable particle size and stiffness (**Figure 1a**). They are comprised of 3.7 wt% PEG backbone, crosslinked via maleimide-thiol linkages, a cell adhesive peptide (RGDS), and a custom macromer for inter-particle annealing (MethMal ^[17]^). Additional PEG-maleimide (PEG-MAL) is incorporated at 0.005 to allow for thiol conjugation. Parallel-step emulsification microfluidic devices were used to generate particles as previously described, using acidic 1x PBS (pH 3.0-4.0) to purify and store particles to slow the hydrolysis of free maleimide groups. Compression testing and fluorescent microscopy was used to analyze formulation stiffness and particle size, yielding particles approximately 90 µm in diameter with a compressive modulus of 26.6 kPa (**Figure 1B-C**).

**Figure 1.**
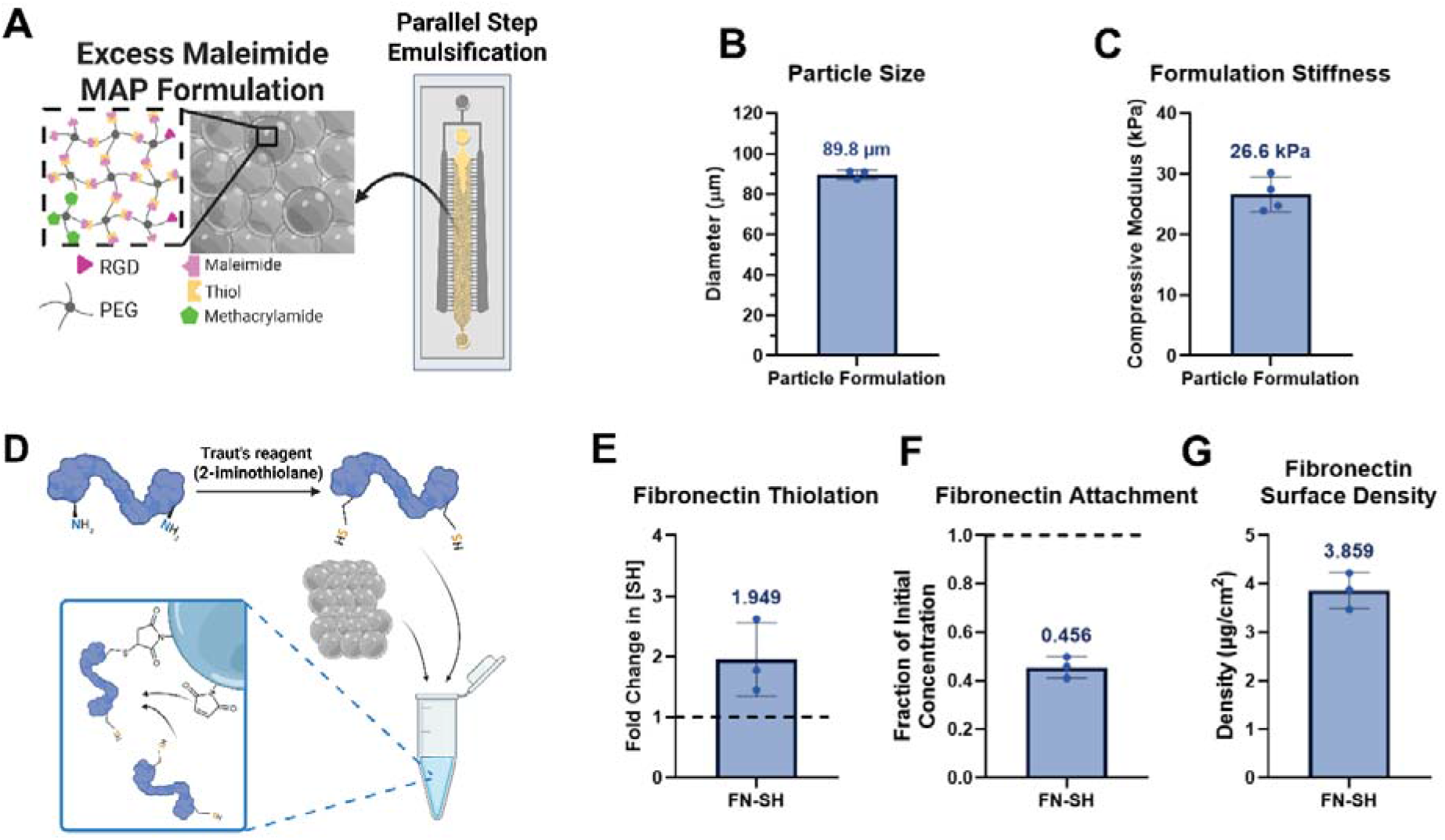
Microparticle synthesis and characterization and protein modification. A) A non-enzymatically degradable formulation was used that relies on Michael-type addition reactions between 4-arm 10kDa PEG-maleimide and PEG-thiol. An RGD cell adhesion peptide and methacrylamide-based annealing macromer are incorporated using the same chemistry, and a stoichiometric excess of maleimide-containing components were incorporated to allow for protein attachment. Particles are synthesized via parallel step emulsification devices that use a fluorosurfactant (Ran Biotechnologies) in NOVEC HFE7500 to form the necessary emulsion. Particles were characterized via fluorescent microscopy for B) size and via compressive mechanical testing for C) stiffness. D) Fibronectin was modified with Traut’s Reagent (2-iminothiolane) for 1 hour at room temperature to convert primary amines to thiols which can then react with the excess maleimide groups on particles to form covalent linkages. E) Fibronectin was successfully thiolated as measured by free thiol concentration compared to an unmodified control. F) Fibronectin was reliably attached to particles as measured by BCA following 20–30-minute incubation at room temperature and the resulting surface density G) was estimated. All graphs show mean +/− standard deviation. All statistics and graph visualization were performed in GraphPad Prism 11. Figure created in part with Biorender (2026).

To facilitate fibronectin attachment to our PEG particles, plasma fibronectin (Gibco) was modified with Traut’s Reagent (2-iminothiolane), which adds to primary amines on lysine residues to yield a free thiol group (**Figure 1D**) and has shown promise for protein modification previously^[34]^. Our particles rely on maleimide-thiol Michael-type additions to crosslink the PEG backbone (Figure 1A) and incorporating additional free thiol groups onto fibronectin via modification of lysine residues allows for conjugation (Figure 1D) while minimally affecting biological activity.^[35]^ Fibronectin was incubated with 10-molar excess of Traut’s reagent and 2-molar excess of Tris(2-carboxyethyl)phosphine (TCEP) to fibronectin in 1x PBS (pH 7.5) with 0.005 EDTA. This solution was purified with Zeba Desalting Columns (7k MWCO, ThermoFisher) and sterilized with 0.22 µm syringe filters before flash freezing for storage. Using a Measure-IT Thiol Assay Kit (Invitrogen), the thiol content on modified fibronectin was measured at 1.95-fold higher than unmodified, indicating successful thiolation (**Figure 1E)**.

Thiolated fibronectin (FN-SH) was incubated with particles containing excess maleimide groups for 30 minutes to tether the protein. Fibronectin can be attached at 45.6% of initial concentration, with an estimated surface density of 3.8 µg cm^-2^ (**Figure 1F-G**). Therefore, the modification was successful and there are sufficient thiol groups to leverage maleimide-thiol chemistry for covalent attachment. This is crucial as it shows FN can be covalently attached to particles and incorporated into the scaffold at surface densities comparable to standard protocols for coating tissue culture surfaces (Roche). These results demonstrate the ability to attach large proteins to granular hydrogels using highly modular click chemistry reactions, which result in surface densities comparable to coatings used in literature. Any changes observed in cell response and mechanical characterization will be due to differences in annealing between these groups, as any modifications were completed after the initial synthesis. Ensuring these relevant properties are kept consistent will minimize confounding effects and isolate the effect of changing protein attachment, as both stiffness and particle size, via its effect on scaffold pore architecture, influence cell behavior.^[12,36]^

### 2.2 Fibronectin-modified MAP scaffolds exhibit increased yield strain and self-healing properties

Following successful attachment of FN to our microparticles, we aimed to characterize the effects of attachment on overall scaffold properties. Scaffold stability (**Figure 2A**) and pore architecture (**Figure 2B**) are both important properties that can impact cell behavior and are highly relevant when considering cell-scaffold interactions. We first wanted to determine the effects on bulk scaffold mechanical properties imparted by the FN attachment, as these properties play an important role in both cell-material interactions and overall scaffold stability. ^[36,37]^ Scaffolds with and without attached FN were tested via parallel-plate rheology with a sandblasted probe and sandpaper-covered plate to prevent slipping at the interface of the scaffold and probe or plate (Figure 2A).

**Figure 2.**
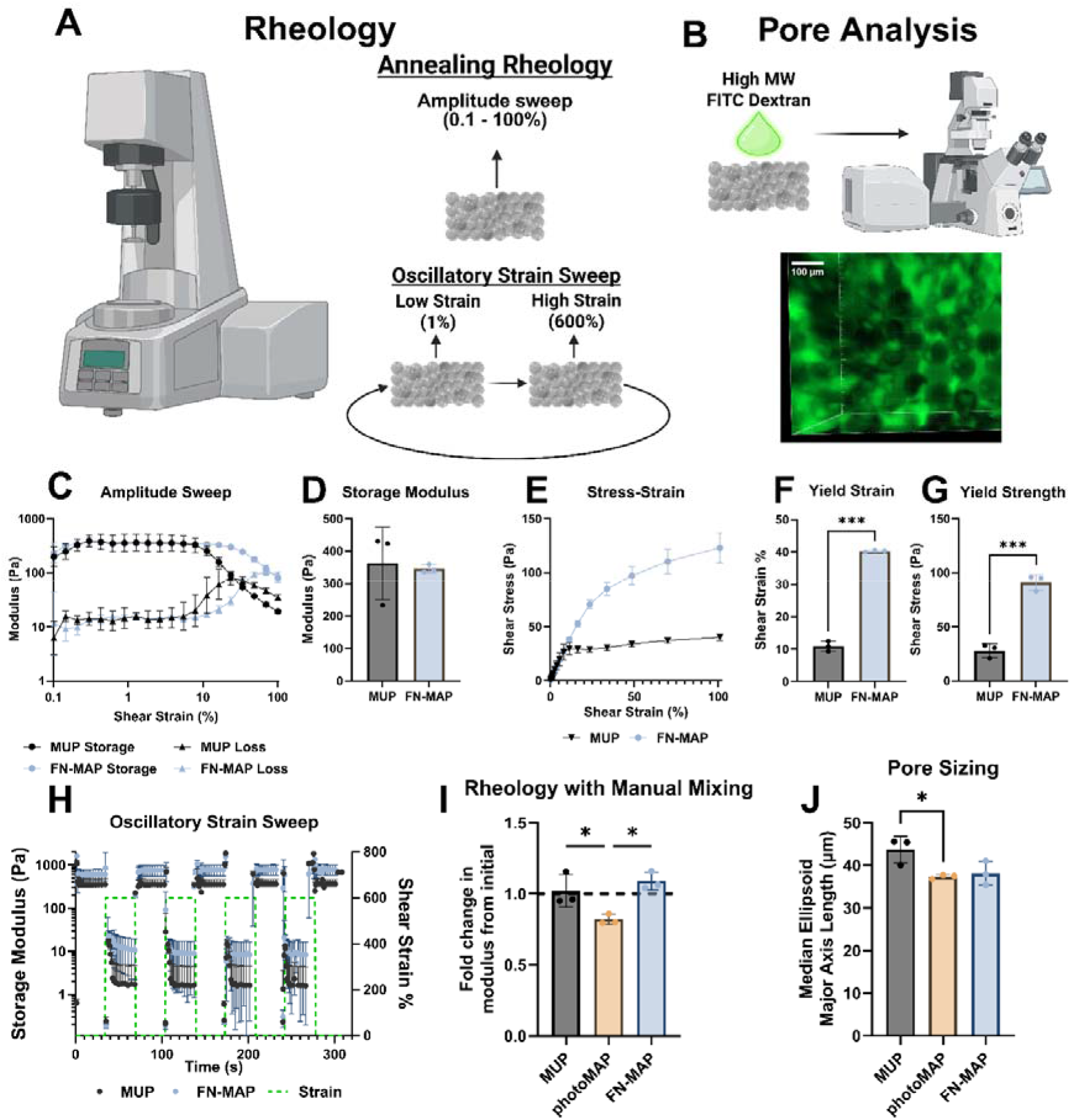
Scaffold characterization via rheology and pore analysis. A) Two rheological tests were performed on scaffolds, including an amplitude sweep to compare MUP and FN-MAP and an alternating strain sweep to determine the self-healing behavior of each group. B) Scaffold pores were visualized by allowing a high-molecular fluorophore-conjugated dextran to diffuse within the pores then imaging using confocal microscopy. C) Amplitude sweeps of MUP and FN-MAP were performed from 0.1 – 100% shear strain with a parallel plate geometry. D) Storage modulus was quantified from the initial flat region of C) and compared with Welch’s t-test. E) Shear stress and shear strain were plotted to obtain stress-strain curves for each group, with the yield strain F) and yield stress G) quantified as described in the methods. Groups were compared with Welch’s t-tests. H) Oscillatory strain sweeps were performed, alternating between 1% strain and 600% strain. I) Scaffolds were also measured before and after manual disruption with a spatula to ensure scaffold destruction and healing. Groups were compared with a one-way ANOVA and Tukey post-hoc testing. J) For each scaffold, the pore architecture was rendered and the shortest ellipsoid axis for each pore was quantified. Medians are shown here. Groups were compared with a one-way ANOVA and Tukey post-hoc testing. *p<0.05; **p<0.01; ***p<0.001; ****p<0.0001. All graphs show mean +/− standard deviation. All statistics and graph visualization were performed in GraphPad Prism 11. Figure created in part with Biorender (2026).

First, an amplitude sweep (**Figure 2C**) was conducted from 0.1% to 100% strain to evaluate the scaffold modulus (**Figure 2D**) and stress-strain relationship (**Figure 2E**). Excitingly, FN-MAP significantly increases the yield stress and strain of scaffolds while not significantly changing the storage modulus of the scaffold, allowing for significantly more stress under rheological shear strain before yielding compared to unannealed samples (**Figure 2F-G**). Notably, these results indicate an ability for FN-MAP to spontaneously self-anneal, in stark contrast to existing MAP systems that often require exogenous crosslinkers or more complex annealing mechanisms.

These results show a clear effect of fibronectin attachment on interparticle scaffold mechanics as it does not affect the overall scaffold stiffness but increases the yield strength by 3-4-fold without exogenous crosslinking, suggesting there are fibronectin-mediated interparticle interactions occurring that stabilize the scaffold in the absence of covalent crosslinking. Particularly, the increased yield strength and strain imply greater strength of these interparticle interactions that allow for increased particle movement before they slip. Verifying these properties gives insight into how FN affects inter-particle mechanics, and this pseudo-crosslinking effect is novel for protein-attached particles. Overall, this result is novel and highly significant as it represents a biologically inspired crosslinking mechanism and shows promise for tissue regeneration applications. With an increased yield strain and stress, these scaffolds can maintain a stable porous geometry under higher strains that might occur in healing tissue like the skin or muscle. In these applications, it would be particularly beneficial for a material to exhibit self-healing and retain strength even after repeated strain on the tissue.

Self-healing is an inherent property of jammed systems, as they can rearrange under strain to retain their bulk structure. However, most covalent annealing methods that increased scaffold mechanics rely on an exogenous stimulus (i.e., light) that does not allow for repeated recovery after strain. Previous iterations of MAP scaffolds have relied on light-based interparticle annealing (photoMAP), which loses mechanical strength after exposure to high strain. To test the self-healing of FN-MAP, we alternated between low and high shear strain with the same setup as above to determine how our material could recover after damage (**Figure 2H**). Additionally, to confirm full scaffold destruction, we tested scaffolds before and after manually mixing and dicing with a spatula (**Figure 2I**).

FN-MAP recovers similarly to MUP, an unannealed granular hydrogel, after repeated periods of high strain (**Figure 2H**). Additionally, FN-MAP can self-heal and retain its mechanical properties after manual disruption of the scaffold where photoMAP fails (**Figure 2I**). These results confirm that this effect is self-healing and reversible; while photoMAP anneals covalently through stable bonds and therefore does not recover after disruption, both MUP and FN-MAP recover after disruption. Finally, MUP is inherently self-recovering, as packed granular hydrogels are, and therefore proves the self-healing ability of FN-MAP while also validating the increased strength of FN-MAP. FN fibril formation is heavily mediated by mechanical forces, as cryptic active sites become exposed under strain that bind to soluble FN and drive fibril assembly.^[38]^ Other possible mechanisms for this increased recoverable strength include disulfide bond formation between adjacent fibronectin monomers, although this is unlikely given there are ≤1 thiols on each protein, or other macromolecular interactions that can reform after strain.

To analyze pore size, scaffolds were formed on glass coverslips and incubated with a high molecular weight (70kDa) dextran tagged with Texas Red (ThermoFisher) to diffuse through the pores over 30 minutes. Fluorescent confocal microscopy was used to image scaffolds over 200 µm and images were analyzed using IMARIS (Oxford Instruments) to create 3D renderings of pores for geometric analysis of median pore ellipsoid length (Figure 2B). FN-MAP was compared to microporous unannealed particles (MUP), a jammed granular PEG hydrogel with no designed interparticle interactions, and photo-initiated microporous annealed particles (photoMAP), a covalently stabilized scaffold using light-mediated radical chain annealing.

FN-MAP allows for stable scaffolds with similar pore sizes to photoMAP and both annealed groups have lower overall porosity than the unannealed group (**Figure 2J**). FN-MAP does not negatively affect scaffold formation and packing compared to photoMAP and aligns with the consistent particle sizes seen in Figure 1F. MUP likely observes increased pore size due to particle movement from adding the fluorescent dye solution. Pore size is a bioactive component of granular scaffolds so ensuring it does not change between crosslinking methods is important as this prevents confounding effects and preserves the essential porosity^[12]^. Because of the difference in pore size, which could become a confounding variable, the MUP group was not investigated in subsequent cellular experiments.

Previous iterations of MAP scaffolds have largely used irreversible crosslinks to maintain structure, while FN-MAP reversibly crosslinks the material while offering higher material strength. Although there are limited examples of reversibly crosslinked granular systems^[18]^, these systems do not include a bioactive linker and often require mixed particle populations or soluble linkers. FN-MAP can provide reversible crosslinking and bioactivity in a homogeneous particle system without requiring an additional soluble component, reducing complexity while adding bioactivity.

### 2.3 FN-MAP allows for higher resolution 3D printing of stable constructs

With these promising rheological findings, we wanted to explore the feasibility of using FN-MAP as a bioink in 3D printing, where the ability of extruded material to maintain structure against body forces after deposition is critical to achieving a 3D structure with the desired feature resolution. Extensive work has been dedicated to designing bioinks and printing processes that yield stable filaments and structures after deposition, and self-healing granular hydrogels make excellent bioinks.^[6,21]^ However, printed particle-based inks, like continuous hydrogels, require additional annealing to maintain longer term structure. Here, we aimed to harness self-healing FN-derived molecular interactions to support natural annealing. Following a previous proof-of-concept study protocol in our group, particles were loaded in a custom syringe and printed at 100 mm min^-1^ (corresponding to an extrusion rate of approximately 70 µL min^-1^) to form a series of parallel lines to assess spreading or a 5-layer square construct to demonstrate stability.^[17]^ Only MUP and FN-MAP were evaluated when printing parallel lines to remove the post-curing step.

After quantifying the thickness of printed lines, we see that FN-MAP allows for higher resolution 3D prints as it resists the characteristic lateral spreading of extruded granular filaments that lack annealing (MUP) (**Figure 3A-B**). FN-MAP allows for rapid self-recovery and stabilization of the extruded structures with inherent self-healing between microparticles preventing loss of structural features over time. Providing higher 3D printing resolution while retaining structure is both excellent evidence for further applications for FN-MAP and corroboration of the rheological results in practice, demonstrating the shear-thinning (printable) and self-healing nature of FN-MAP. FN-MAP also provides sufficient strength for printed constructs to retain their structure through agitation (**Figure 3C**) similarly to covalently annealed constructs (photoMAP). FN-MAP is shear-thinning but can self-heal back to its original strength, allowing for prints with comparable strength to covalent linkages without requiring exogenous crosslinking. FN-MAP can be 3D printed effectively and retain its structure, offering an excellent choice for an innately bioactive, self-healing, shear-thinning 3D bioink that requires no exogenous crosslinking.

**Figure 3.**
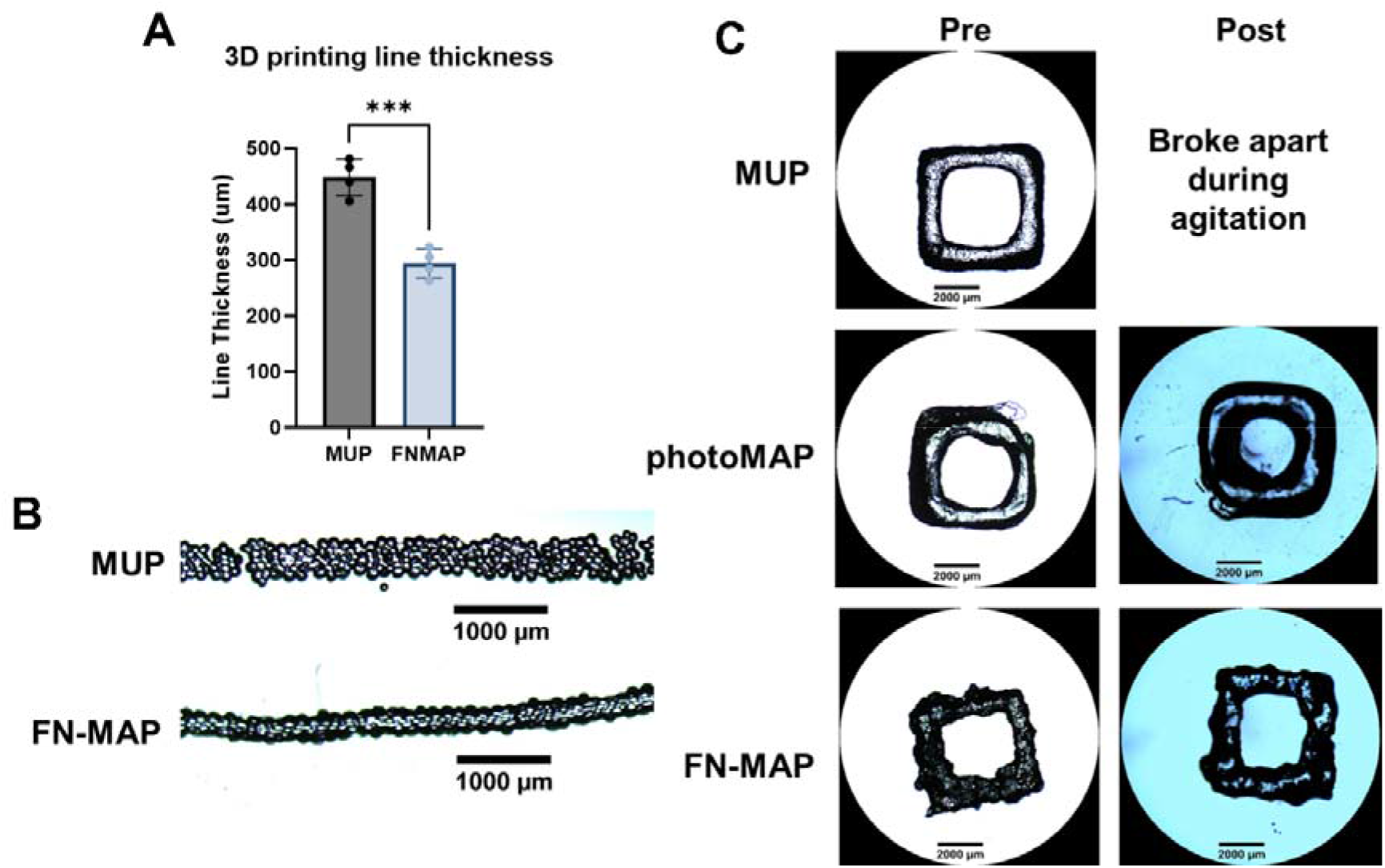
Application of FN-MAP in 3D printing. MUP and FN-MAP were loaded into custom syringes for extrusion-based 3D printing, as described in 4.4. A-B) 4 single-layer lines were printed for each group and the thickness of each line was measured at 5 points. Groups were compared with a Welch’s t-test. C) 5-layer squares were printed (and, for MAP, exposed to 365 nm light to anneal), then imaged before and after submerging in PBS with shaking for 5 minutes. *p<0.05; **p<0.01; ***p<0.001; ****p<0.0001. All graphs show mean +/− standard deviation. All statistics and graph visualization were performed in GraphPad Prism 11. Figure created in part with Biorender (2026).

### 2.4 FN-MAP encourages cell spreading and facilitates more active fibroblast phenotypes

In addition to the significance of its inherent material properties and stability, our hypothesis for FN-MAP also focused on its bioactivity, corresponding to the ability of FN to influence fibroblast behavior via endogenous ECM signaling. Normal human lung fibroblasts (nhLF) were used for their widespread availability and relevance across a range of clinically important applications, including pulmonary disease modeling, tissue reconstruction, and wound healing research.^[39,40]^ Fibroblast spreading is an indicator of cell-matrix interactions and functional activation state. Fibroblasts bind fibronectin primarily via α5β1 integrins through the RGD motif, and this interaction signals through canonical mechanotransduction pathways (including RhoA/ROCK and YAP/TAZ) to yield the characteristic elongated, contractile morphology associated with an active fibroblast phenotype.^[41–43]^ Therefore, fibroblast morphology can be a helpful indicator for whether the tethered FN is presenting accessible binding sites and eliciting downstream signaling. Importantly, a more spread and elongated morphology can be associated with a more functional, less quiescent fibroblast state. A prominent characteristic of fibroblasts in aging tissue is reduced spreading and contact with ECM, resulting in a collapsed morphology as cells lose their typical elongated spindle-like morphology and become shorter, rounded, and collapsed.^[44]^ Promoting spread, elongated morphologies is therefore desirable for tissue regeneration, as it is associated with productive matrix engagement rather than a quiescent state without necessarily driving pathological fibrosis, which requires sustained activation beyond what is expected in this context.

Cells were seeded within the interstitial space of photoMAP or FN-MAP scaffolds to remove any differential effect of cell migration. After 48 hours of culture, scaffolds were fixed and stained to visualize nuclei and cytoplasm and analyzed to evaluate cell morphology in Imaris (**Figure 4A-B**). Compared to photoMAP, FN-MAP increases cell cytoplasm area, elongation length (ellipsoid axis length), and decreases sphericity, all indicating that it encourages fibroblasts to spread and adopt more elongated morphologies (**Figure 4C-E**). In addition to our existing results on the mechanical effects of fibronectin attachment, this data provides evidence for additional bioactivity imparted by the FN. Cells are more likely to spread and interact with particles in FN-MAP compared to photoMAP, suggesting that cells are meaningfully engaging with tethered FN and these interactions are leading to downstream morphological changes that could increase ECM production.^[45,46]^ These results are encouraging, as they demonstrate FN can retain at least some of its inherent bioactivity even while tethered to particles, although the complete effects on downstream fibroblast function and behavior were not investigated. Additional characterization is needed to understand how covalently tethered FN presents to cells, which limits its viscoelastic behavior and may impact binding affinity.

**Figure 4.**
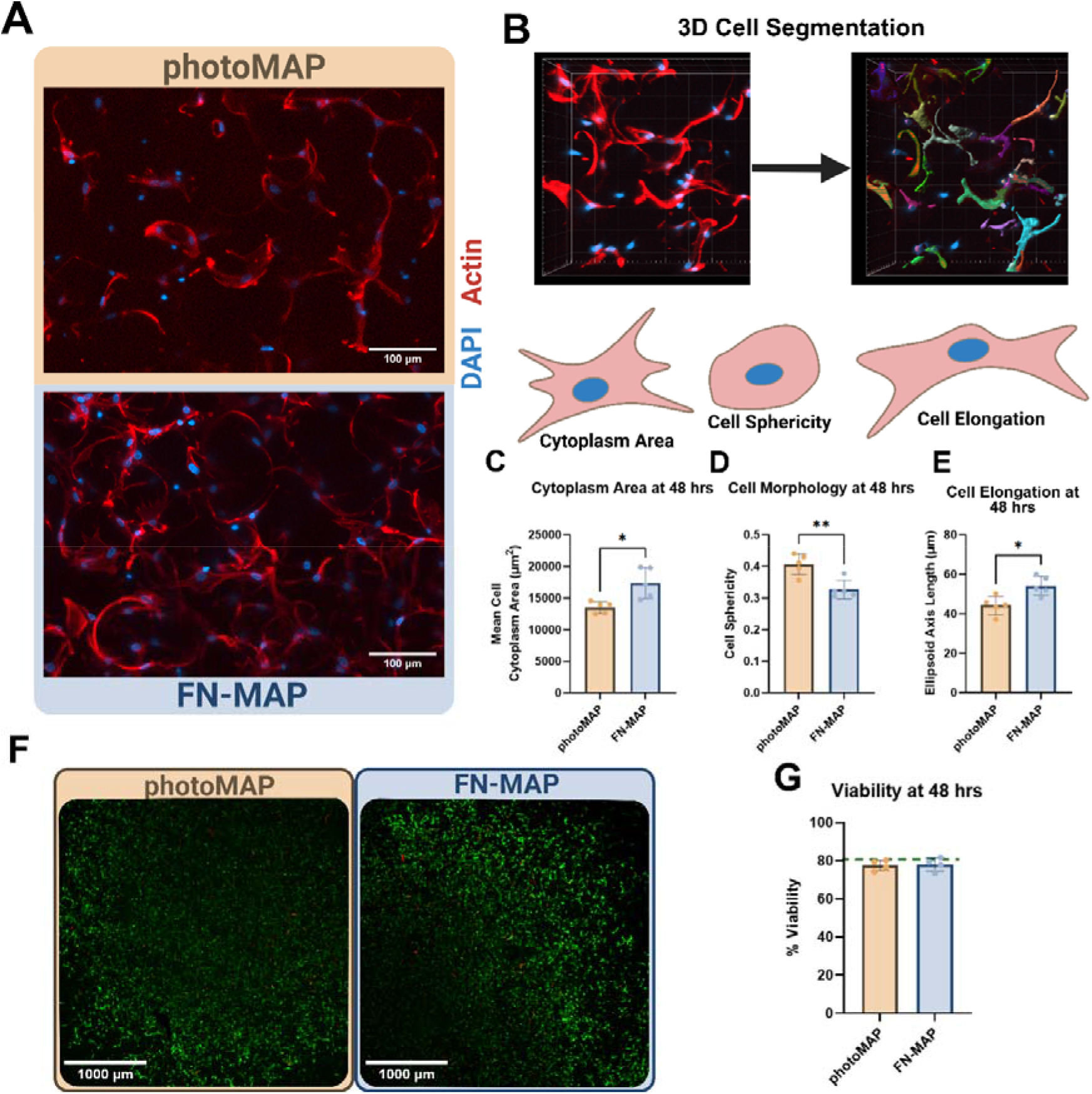
Cell morphology and viability in response to FN attachment and annealing chemistry. A) Representative maximum intensity projection (MIP) images from each group showing human lung fibroblasts (HLFs) and their nuclei (blue, DAPI), actin (red, phalloidin-647) after 48 hours of culture. Scale bars represent 100 µm. B) Z-stacks were processed via 3D cell segmentation in IMARIS to analyze cell shape and morphology. C-E) HLF morphology metrics including cytoplasm area C), sphericity D), and elongation E). F-G) HLF viability after 48 hours of culture in each group, showing a high percentage of live (green) cells compared to dead (red) cells). All groups in C)-E) and G) were compared with Welch’s t-test. *p<0.05; **p<0.01; ***p<0.001; ****p<0.0001. All graphs show mean +/− standard deviation. All statistics and graph visualization were performed in GraphPad Prism 11. Figure created in part with Biorender (2026).

Given the observed rheological properties of FN-MAP, we wanted to ensure that the changed annealing behavior provided by FN did not adversely affect cell viability. Following 24 hours of culture after seeding, FN-MAP shows no detrimental effect on cell viability compared to covalently annealed photoMAP (**Figure 4F-G**). Ensuring the viability is maintained between groups is significant and supports our hypothesis that any changes in cell behavior will be due to fibronectin incorporation and validates that the FN-MAP strategy is non-cytotoxic.

### 2.5 FN-MAP accelerates epidermal closure while allowing for cell infiltration

To evaluate whether the beneficial effects on fibroblast morphology imparted by FN-MAP would translate to *in vivo* applications, we turned to a model of impaired tissue regeneration in diabetic chronic wound healing. In the normal wound healing process, a cascade of signals orchestrates tissue repair and regeneration.^[47–49]^ Neutrophils and macrophages infiltrate early on to clear the site of pathogens, navigating via fibrin-based ECM, while keratinocytes and endothelial cells migrate to repair the fragile epidermis and vasculature, respectively. This nascent ECM gives way to granulation tissue and, eventually, more mature tissue, as fibroblasts migrate and begin secreting collagens and proteoglycans.

In diabetic patients, this process is significantly disrupted in a variety of ways, including impaired vascularization and epithelial regeneration, and has been an excellent candidate for biomaterial scaffolds.^[50,51]^ We sought to use a diabetic mouse model to confirm that the increased mechanical properties and bioactivities of FN-MAP would translate well to a strained wound healing environment. Modifying the protocol from Pruett et al.,^[4]^ mice were wounded under anesthesia and treated with photoMAP or FN-MAP covered with silicon scarsheets (**Figure 5A**). Wounds were harvested after three days and embedded for sectioning and staining. From gross H&E images, the scaffolds appear similar, experiencing limited cell infiltration but remaining intact, despite annealing via different mechanisms (**Figure 5B**).

**Figure 5.**
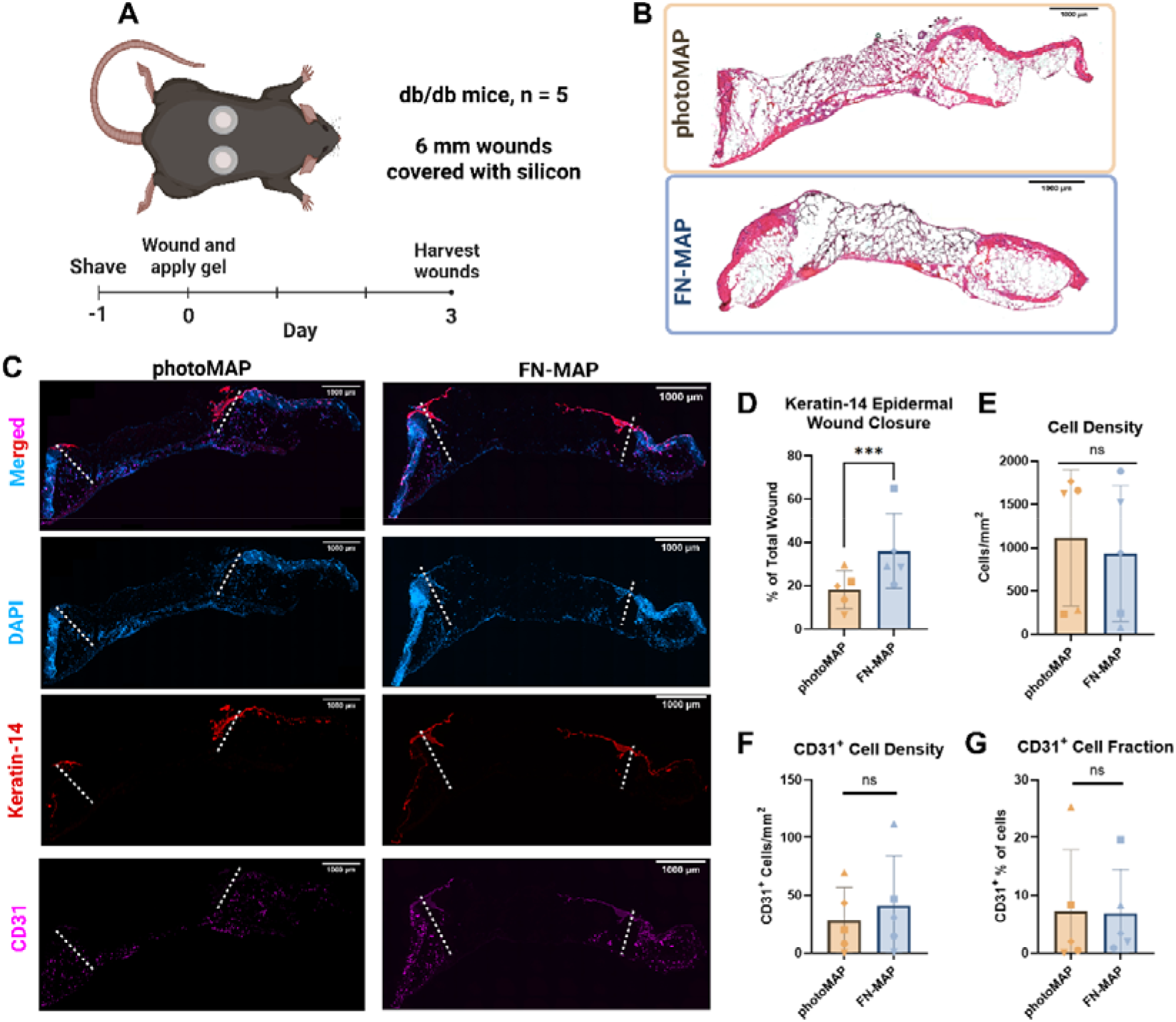
FN-MAP as a scaffold for wound healing. A) Male diabetic mice (db/db) were first shaved and had their nails clipped, then were wounded under anesthesia with a 6 mm biopsy punch. Wounds were covered with silicon scarsheets to prevent drying out and were harvested after sacrifice at day 3. B) Representative images of H&E staining on sections from wounds from each group. Scale bars represent 1 mm. C) Representative fluorescent images of wounds stained with DAPI (blue), Keratin-14 (red), and CD31 (magenta). Scale bars represent 1 mm. Wound boundaries are marked with white dashed lines. D) Epidermal closure was quantified as the ratio of keratin-14 positive epidermal length to the total wound surface length. E) Total cell density and F) CD31+ cell density were quantified, along with G) CD31+ cell fraction, defined as the percentage of total cells positive for CD31. For D-G), each mouse is denoted with a unique symbol. Statistical results for D-G) were derived from linear mixed models using the lmer and emmeans packages in R to account for mouse-to-mouse variation and wound location variation. *p<0.05; **p<0.01; ***p<0.001; ****p<0.0001. All statistics and graph visualization were performed in GraphPad Prism 11 or R (4.3.2). Figure created in part with Biorender (2026).

After immunofluorescent staining with keratin-14 (**Figure 5C**), we observed that FN-MAP increases wound closure as measured by epidermal tongue length normalized to wound length, indicating it can facilitate faster wound closure than photoMAP (**Figure 5D**). These results are promising, as wound closure is a highly relevant metric for wound healing. This increase in epidermal closure could be due to a combination of the bioactive effects on cell spreading in vitro and the high mechanical integrity of FN-MAP scaffolds, allowing for more stable scaffolds that allow for increased cell spreading. Wound closure rate is a highly clinically relevant metric for wound healing, as faster epidermal closure helps to prevent infection and allow for more complete healing within the tissue. Accelerating this process is very promising for application of FN-MAP scaffolds and demonstrates that these results go beyond scaffold architecture alone; therefore, the bioactive effects observed *in vitro* are likely translating to the wound environment.

In contrast, when investigating endothelial cell migration (CD31^+^) and overall cell infiltration (Figure 5C), there were no significant differences observed, although there was a significant amount of variability in these samples (**Figure 5E-G**). Samples from the same mouse are denoted by the same symbols in Figures 5D-G and were evaluated with pairwise comparisons derived from a linear mixed model accounting for mouse-to-mouse and wound location variability, as samples appeared to exhibit similar behavior across different metrics.. These results are unexpected given the effect on cell spreading and adhesion seen previously, but given the highly inflammatory environment, it is possible the cell-material interactions were dampened. Additionally, in this environment, the material may not attract cells to the same degree as it requires physical cell-material interactions to affect behavior.

Vascularization is critical to long-term scaffold integration and tissue regeneration, although previous groups have shown success with MAP scaffolds^[4,5]^. The lack of a significant difference in CD31^+^ infiltration or total cell density may be related to the complex wound environment, where host-derived fibronectin and competing chemotactic signals may dilute scaffold-specific effects. However, in contrast to other infiltrating cell types that migrate in 3D, keratinocytes migrate primarily in 2D across the surface of the scaffold in the wound bed, decreasing their distance to travel. These other cell types, including fibroblasts, macrophages, and endothelial cells, may be affected more significantly by the complex wound environment in their ability to respond to the scaffold. The bioactive effects of FN-MAP were observed *in vitro* on fibroblast behavior, while macrophages and neutrophils are the first responders in wound environments.^[47,52]^ Given these results were observed in a diabetic wound environment, which often experiences significantly impaired angiogenesis,^[4,51]^ future iterations may require a chemoattractant or other signaling molecules to affect blood vessel infiltration.

## 3. Conclusions

Although a variety of strategies exist to incorporate additional bioactivity and stability into synthetic granular hydrogels, this manuscript describes a novel approach wherein we coat synthetic granular hydrogels with fibronectin (FN-MAP), resulting in a scaffold capable of ECM-inspired dynamic annealing. We demonstrate increased mechanical stability via rheology and increased bioactivity *in vitro* and *in vivo* compared to RGD-coated light-annealed granular hydrogels (photoMAP). Under repeated high strain, FN-MAP recovers and maintains its stability while photoMAP is unable to remain stable; additionally, FN-MAP exhibits significantly increased yield-strain and strength, indicating the presence of interparticle interactions. The self-healing ability of FN-MAP allows for promise as 3D printing bioink, providing higher resolution and more stable prints. Analysis of cell interactions *in vitro* shows increased cell spreading and comparable viability of FN-MAP to photoMAP, and FN-MAP accelerates epidermal closure in a diabetic wound healing model. These results begin to demonstrate its ability to influence cell movement and cell-material interactions in beneficial ways while increasing scaffold strength. Altogether, we present a novel biomaterial that offers increased stability and bioactivity without mixed particle populations or exogenous linkers, providing consistent results using a biologically inspired design. Future directions will explore the mechanism of interparticle interactions, potentially leveraging Fluorescence Resonance Energy Transfer (FRET) microscopy to characterize these interactions. Additional applications will also be explored for tissue regeneration and repair in clinically relevant models including tissue bulking for vocal folds and cell delivery.

## 4. Experimental Methods

### 4.1 Fibronectin modification

#### 4.1.1 Materials for modification

Plasma fibronectin (Gibco) was dissolved at 2 mg mL^-1^ in sterile 1x phosphate-buffered saline (PBS), in accordance with manufacturer’s guidelines. Traut’s reagent (2-iminothiolane, ThermoFisher) was dissolved at 2 mg mL^-1^ in ultrapure (MilliQ) water, Tris(2-carboxyethyl) phosphine (TCEP, ThermoFisher) was dissolved at 0.01 in 1x PBS, and ethylenediaminetetraacetic acid (EDTA, Sigma-Aldrich) was dissolved at 0.2 in ultrapure (MilliQ) water. All solutions were prepared immediately prior to use.

#### 4.1.2 Thiolation and Sterilization

To thiolate the fibronectin, Traut’s reagent was added at a 10-molar excess per mol fibronectin, TCEP was added at a 2-molar excess per mol fibronectin, and EDTA was added to achieve a final concentration of 0.002 M. Once combined in a low-binding tube, the solution was allowed to react on a tube rotator for 1 hr at room temperature. Zeba Desalting Columns (7k MWCO, 0.5 mL, ThermoFisher) were used to purify the protein solution according to manufacturer instructions. The solution was then diluted to 0.44 mg mL^-1^ in 1x PBS and additional TCEP (2-molar excess per mol fibronectin) was added to prevent disulfide bond formation. This final solution was sterilized via syringe filtration through a 0.22 µm filter (VWR) and aliquoted into low-binding tubes before flash freezing (stored at -20°C short-term, -80°C long-term). A Measure-IT Thiol Assay Kit (ThermoFisher Invitrogen) was used to quantify the thiol concentration on thiolated and unmodified fibronectin (both treated similarly with TCEP and EDTA) according to manufacturer instructions.

### 4.2 Microparticle fabrication

#### 4.2.1. Materials and formulation

4-arm poly(ethylene glycol) (PEG) maleimide (PEGMAL, 10kDa and 20kDa) and 4-arm PEG-thiol (PEG-SH, 10kDa) were purchased from Nippon Oil Foundry (Japan), RGD cell adhesive peptide (Ac-RGDSPGGC-NH_2_) was purchased from WatsonBio, and reagents involved in the synthesis of a custom annealing macromer (MethMal, 4-arm PEG-maleimide modified to have 2-3 methacrylamide-terminated arms) were purchased from Sigma-Aldrich. MethMal synthesis was performed as previously described and allows internal crosslinking of particles to be decoupled from inter-particle annealing.^[17]^

Particles were synthesized at 3.7% (w/v) PEG with excess PEGMAL (0.005) to ensure unreacted maleimides were available for conjugation. Final concentrations for all components were: 38.533 mg mL^-1^PEGMAL, 0.001 RGD, 0.001 MethMal, 5 x 10^-6^ biotin-maleimide for streptavidin-fluorophore visualization, and 18.252 mg mL^-1^ PEG-SH.

#### 4.2.2. Microparticle generation

Parallel step emulsification microfluidic devices^[53]^ with a 19.7 µm channel height were used to fabricate particles following existing protocols^[54]^. Briefly, devices were casted with polydimethylsiloxane (PDMS) from master molds, bonded to glass slides, and treated with a (trichloro(1H,1H,2H,2H-perfluorooctyl) silane) solution (1:50). PEGMAL, RGD, and MethMal were dissolved sequentially in 10x PBS (pH 1.9), and PEG-SH and biotin-maleimide were dissolved in 1x PBS (pH 7.4). These solutions were combined 1:1, loaded into syringes, and flowed through the microfluidic devices with a fluorosurfactant (2% w/v, Ran Biotech) diluted in NOVEC-7500 (3M) at a 2:1 flow rate (oil: aqueous) controlled via syringe pump. Particles were crosslinked overnight at room temperature, purified with serial washes (and centrifugation) and transitions in 1x PBS (pH 3-4), and sterilized in 70% IPA before storage at 4°C.

#### 4.2.3. Microparticle characterization

##### Sizing

Particles were incubated with DyLight 650-conjugated streptavidin (ThermoFisher) to fluorescently label them via high affinity biotin-streptavidin binding. The particles were imaged via spinning disk confocal microscopy (Molecular Devices) and analyzed using a custom ImageXpress macro to quantify particle areas used to calculate particle diameter and polydispersity index (PDI). Particles from every batch were imaged and analyzed to ensure consistent sizes across batches.

##### Stiffness

To estimate particle stiffness, the same formulation used to generate particles was used to create macrogel cylinders for bulk compression testing. 0.1 mL of the gel solution was pipetted on slides coated with Sigmacote© (to prevent adhesion) and lined with 2 mm silicon spacers. The macrogels were allowed to react overnight, then placed in 1x PBS to swell. Macrogels were tested with a 5N load cell on an Instron mechanical tester (BlueHill) with a strain rate of 0.5 mm min^-1^. A custom Matlab (Mathworks, R2023b) script was used to analyze the slope of the resultant stress-strain curve between 10 and 20% strain to quantify the compressive modulus.

#### 4.2.4. Fibronectin attachment

##### Attachment

Attachment must be performed with 24 hr of synthesis to ensure that enough maleimide groups are available for conjugation, as maleimides hydrolyze rapidly. To attach fibronectin to the particles, FN-SH (0.44 mg mL^-1^) was incubated with particles at a 1-1.5:1 ratio (v/v) for 20-30 minutes at room temperature on a plate shaker or tube rotator. After incubation, the solution was centrifuged at 18000 x g for 5 minutes and the supernatant was removed to isolate the particles. Particles were stored in sterile 1x PBS at 4°C for up to 1 week before use. Particles without FN for photoMAP were incubated with 0.002 lithium phenyl-2,4,6-trimethylphosphinate (LAP) at 10% of the particle volume in 1x PBS for at least 15 min.

##### Characterization

To quantify the FN concentration in the particle slurry, a bicinchoninic acid (BCA) assay was run on particles incubated with and without FN-SH according to manufacturer protocol. A standard curve of FN-SH diluted in 1x PBS was used to interpolate the concentration in the particles, and absorbance at 562 nm was read using a plate reader (Biotek).

### 4.3 Scaffold characterization

#### 4.3.1 Rheology

##### Material Preparation

Particles with and without attached FN were isolated via centrifugation (18,000g for 5 min) and supernatant aspiration. Particles without FN were split into two groups (MUP and photoMAP), one with LAP photoinitiator (photoMAP) and one without (MUP). After incubation with 1x PBS or LAP, particles were isolated as above in preparation for rheological measurements. An Anton Paar Modular Compact Rheometer (MCR 302) was used to acquire all rheological data in this study. Particles were centrifuged twice at 21,000g for 5 minutes to approximate consistent packing.

###### Rheological Measurements – Amplitude Sweep

55 µL of particles were pipetted on the center of a UV-compatible stage. A UV lamp was calibrated to yield 33.4 mW cm^-2^ on the scaffold, as measured via spectrophotometer at 365 nm. An 8 mm sandblasted parallel plate probe was lowered to 0.9 mm above the plate to ensure contact with the scaffold. A prestress step (20% strain, 4 Hz) was performed to remove deformities in the scaffold. All samples were exposed to UV light for 29 seconds, although only photoMAP samples contained photoinitiator. Samples were then tested with an amplitude sweep from 0.1% -100% strain at a constant frequency (1 Hz). Three independent samples (n = 3) were tested for each condition. The data was exported to Microsoft Excel and analyzed with a custom script in Python to identify the storage modulus, denoted as the constant region of Figure 2C, and the yield strength, denoted as the end of the linear region in the stress-strain curve in Figure 2D. Yield strain was interpolated using a smoothing spline with 5 knots to approximate strain at the yield strength.

###### Rheological Measurements – Oscillatory Strain and Manual Mixing

55 µL of particles were pipetted on the center of a UV-compatible stage. A UV lamp was calibrated to yield 33.4 mW cm^-2^ on the scaffold, as measured via photospectrometer at 365 nm. An 8 mm sandblasted parallel plate probe was lowered to 0.9 mm above the plate to ensure contact with the scaffold. An oscillatory strain sweep was performed, alternating between 1% (low) and 600% strain to determine the recovery behavior of FN-MAP, with a total of 4 completed cycles. Additional samples were tested for a single cycle at 1% strain, manually mixed with a spatula to ensure the entire scaffold was disrupted, and tested again under the same conditions. The fold change from the initial measurement of the storage modulus, calculated by average 5 points in the middle of each cycle to eliminate effects due to strain changes, was calculated for each sample.

#### 4.3.2 Pore sizing

Analysis of scaffold pore architecture was performed as previously described.^[9]^ Briefly, particles prepared as above in the section “Material Preparation” for 4.3.1 were transferred into custom scaffold molds make by punching 3 mm holes in DuoDerm adhered to a glass coverslip. All samples were exposed to UV light (33.4 mW cm^-2^ for 29 seconds, 365 nm) then incubated with a solution of 2 MDa FITC-Dextran for 1-2 hr in a dark, humidified chamber. The high molecular weight prevents diffusion into particles and restricts fluorescence to the pore space. A Zeiss LSM980 laser-scanning confocal microscope (Advanced Microscopy Facility at the University of Virginia, RRID: SCR_018736) was used to image scaffolds at two sites in each scaffold. Z-stack images were obtained with a 2 µm step size (minimum of 100 steps). Images were analyzed in IMARIS (Oxford Instruments) by first performing deconvolution (with the appropriate microscope-specific parameters) and a Gaussian filter and subsequently rendering surfaces to match the pore space. Seed points were automatically generated to morphologically segment individual pores before exporting statistical parameters. Specifically, ellipsoid axes are calculated for each pore; selecting the smallest axis length approximates the diameter of the largest sphere that can fit in the pore. The median axis length for each site was calculated and averaged within a single sample.

### 4.4 3D Printing

#### 4.4.1 Materials and Printer

Particles were prepared as in the *Material Preparation* section of 4.3.1 and loaded into a syringe. 3D printing was performed using a FELIX 3.1 3D printer (FELIXprinters, IJsselstein, Netherlands) whose printhead was modified for positive-displacement extrusion from a syringe, according to a published design.^[55]^ Repetier-Host software (Repetier) was used to control extrusion guided by custom G-codes. A translation speed of 100 mm min^-1^ was used for printing which, given the feed rates specified in the G-code used, correlated to an approximate volumetric extrusion rate of 70 µL min^-1^ or, equivalently, 7 µL of particles per 10 mm of translation.

#### 4.4.2 Line Printing

For each condition, 3-4 lines were extruded (at least 5 mm in length) on the bottom of a dish containing a wet Kimwipe to provide moisture and prevent excess drying. After printing, dishes were sealed with parafilm then imaged on a widefield microscope (EVOS, ThermoFisher) in brightfield. 2 regions for each line were analyzed for width in ImageJ via manual measurements.

#### 4.4.3 Construct Printing

For each condition, 5-layer squares were printed (each 1 mm x 1 mm) into humidified petri dishes as in the previous section. Samples were exposed to UV light (33.4 mW cm^-2^ for 29 seconds) then imaged on a widefield microscope (EVOS, ThermoFisher). 1x PBS was carefully added to the dish and the dishes were placed on shaker plate for 5 minutes at 100 rpm. After agitation, the squares were imaged again (if possible).

### 4.5 In vitro experiments

#### 4.5.1. Materials and general methods Materials

Particles were prepared and sterilized as described in 4.2. They were spun at 18,000 x g for 5 minutes to dry before proceeding with cell experiments. Normal human lung fibroblasts (HLFs) were purchased from ATCC. For *in vitro* experiments, HLFs were thawed and cultured according to manufacturer protocol till confluency. All cells were used at either passage 5 or 6 to ensure consistency.

#### 4.5.2. Cell spreading assay

HLFs were passaged according to manufacturer protocol. Cells were pelleted to achieve a density of 2000 cells µL^-1^ of material, aspirated then dried particles were added and mixed thoroughly. 25 µL scaffolds were plated in treated 96-well plates, incubated under standard HLF conditions for 48 hours, then washed with 1x PBS and fixed with 4% paraformaldehyde (PFA) for 15 minutes before storage in 1x PBS at 4°C. These scaffolds were blocked with 100 µL 3% (w/v) bovine serum albumin (BSA) in PBS-T (0.3% (v/v) Triton-X in 1x PBS), then stained for actin using phalloidin-AlexaFluor 647 at 1:200 (Invitrogen) and DAPI (Invitrogen) at 1:1000 overnight at 4°C. Scaffolds were imaged with a Zeiss LSM980 laser-scanning confocal microscope (Advanced Microscopy Facility at the University of Virginia, RRID: SCR_018736) using a 20x objective at 1.1 µm step sizes over 150 µm at multiple sites per scaffold. The raw .czi files were analyzed in IMARIS (Oxford Instruments) to create 3D surfaces of cell bodies and nuclei. A batch process was run to deconvolute the images and smooth them, then each image was analyzed to generate surfaces. Statistics about cell morphology and position were exported and organized in R 4.3.2 then analyzed and visualized in GraphPad Prism.

#### 4.5.3 Cell viability

Cells were seeded as described in 4.5.2, with an increased cell density of 5000 cells µL^-1^. After 24 hours, scaffolds had their media removed and were washed with sterile 1x PBS. The Live/Dead Viability Kit (Invitrogen) was prepared according to manufacturer protocol and added in equal volume to each scaffold before incubating for 20 minutes. Imaging was performed with a Molecular Devices ImageXpress MicroConfocal at 10x magnification with 50 µm step sizes (for 400-500 µm). Analysis was done in FIJI (NIH) using the “Analyze Particles” function and viability was quantified as the fraction of total cells marked as live.

### 4.6 Wound healing assay

#### 4.6.1 Mouse Wound Healing Model

All surgeries were performed in accordance with Animal Care and Use Committee guidelines under UVA Animal Protocol #4165. 16-week-old male db/db mice (Jackson stock: 000697) were used for the study. Five mice were evaluated on Day 3.

#### 4.6.2 Mouse Wound Healing Protocol

Mice were anesthetized using continuous administration of isoflurane. Hair was removed from the dorsal side of the mouse using electric clippers and Nair, followed by a PBS wash. Nails were trimmed to reduce the likelihood of splint removal. Before surgery, mouse backs were sterilized with iodine and 70% isopropanol. A sterile 5 mm biopsy punch was used to mark up to four wound areas, followed by cutting the wound area using surgical scissors. Four wound sites were included to allow for an unannealed (MUP) and clinical (Oasis) treatment group, but these two groups were excluded due to difficulties in processing and instability. Each treatment was applied to a wound in a randomized order, rotated for each mouse. 15 µL of each MAP scaffold condition was applied and smoothed on the surface of the wound using a positive displacement pipette. photoMAP scaffolds were annealed for 30 seconds using 365 nm UV light (15.4 cm away, 440 mA, Thor Labs). Silicone (ScarAway) circles were used as wound coverings and splints. Prior to surgery, two silicone patches were stacked on top of each other and then cut using a 12 mm biopsy punch. The silicone splints were superglued (Gorilla Glue) around the wound to prevent contraction. Mice received 0.1 cc of Buprenorphine immediately following surgery, were singly housed, and checked twice a day to confirm they had not removed their bandages. Mice were sacrificed with an isoflurane overdose and cervical dislocation after three days. The wound samples were removed using surgical scissors and immediately embedded in OCT on dry ice for later cyro-sectioning.

#### 4.6.3. Tissue processing and staining

OCT blocks were sectioned on to positively charged glass slides (Fisherbrand) at 14-18 µm thickness with at least three section on each slide. Slides were allowed to come to room temperature, then covered with 4% paraformaldehyde for 15 minutes and subsequently rinsed 3x in 0.3% Triton-X in PBS (PBS-T). They were blocked in 5% goat serum (diluted in PBS-T) for 30 minutes at room temperature, then incubated with primary antibodies diluted in 2% goat serum in PBS-T overnight at 4°C. Antibodies used in this study were for Keratin-14 (Biolegend 905304, rabbit) at 1:100 and CD31 (BD 553370, rat) at 1:150 (both v/v). Slides were washed in PBS-T and incubated with secondary antibodies (Goat anti-rat AlexaFluor 488 and goat anti-rabbit AlexaFluor 647) and DAPI (1:1000) for 2 hours at room temperature before washing in PBS and mounting using Prolong Anti-Fade Gold (Invitrogen). All stained slides were stored at 4°C in flat folders until imaging.

#### 4.6.4. Imaging and Analysis

A Zeiss AxioScan 7 was used to scan all slides in this study with a 20x objective. QuPath v0.6.0^[56]^ was used to analyze images. Briefly, all implants were annotated and visually checked to ensure quality. To analyze blood vessel infiltration, an object classifier was trained using at least one image from each group and run on the implants to yield values for both general cell infiltration (using DAPI) and CD31^+^ cell infiltration (using CD31 and DAPI). Epidermal closure was also quantified by tracing the total wound length along the surface and tracing the epidermal tongue growth, then normalizing epidermal growth to wound length. Analysis was performed as described in 4.7.

### 4.7 Statistical Analysis

All statistical tests and visualizations were done in GraphPad Prism 11. All analyses are noted in figure captions or described in the corresponding methods section. P-values greater than 0.05 were considered significant. Graphs are represented with mean ± standard deviation. *p<0.05; **p<0.01; ***p<0.001; ****p<0.0001. For analysis of epidermal closure and cell infiltration in 4.6, we used the *lmer*^[57]^ package in R to create linear mixed models with random effects associated with mouse and wound location variability and fixed effects due to scaffold group. Using *emmeans*, we ran pairwise comparisons between scaffold groups to isolate these effects.

## Acknowledgements

The authors would like to thank the National Institute of Diabetes and Digestive and Kidney Diseases (NIDDK) (5 R01 DK126020) and the National Institute on Deafness and Other Communication Disorders (NIDCD) (5 R01 DC020998). The content is solely the responsibility of the authors and does not necessarily represent the official views of the National Institutes of Health. We would also like to thank the Advanced Microscopy Core at UVa, particularly Dr. Sijie Hao, for his valuable guidance on confocal microscopy. This work used the Zeiss LSM980 in the Advanced Microscopy Facility which is supported by the University of Virginia School of Medicine (RRID): SCR_018736. Figure schematics were developed with Biorender (2026) and GraphPad Prism 11.

## Data Availability Statement

((include as appropriate, including link to repository))

Received: ((will be filled in by the editorial staff))

Revised: ((will be filled in by the editorial staff))

Published online: ((will be filled in by the editorial staff))

